# Expression of GDNF Receptors Within Nucleus Ambiguus During Rat Development

**DOI:** 10.1101/2022.05.04.490634

**Authors:** Quinton Blount, Ignacio Hernandez-Morato, Yalda Moayedi, Michael J Pitman

**Affiliations:** Mercer University School of Medicine, Columbus, GA; Department of Otolaryngology-Head & Neck Surgery, Columbia University College of Physicians and Surgeons, New York, NY; Department of Neurology, Columbia University College of Physicians and Surgeons, New York, NY

**Keywords:** Laryngeal proprioception, muscle spindles, VGLUT1, intralaryngeal ganglia, dysphonia, nucleus ambiguus, GDNF, Ret, GFRalpha, larynx, recurrent laryngeal nerve

## Abstract

**Objective:** Upregulation of GDNF and its receptors is observed during laryngeal reinnervation after nerve injury. In contrast, little is known regarding the expression of GDNF receptors in the formation of the nucleus ambiguus (NA) and its innervation of the larynx during embryogenesis. Differences may suggest therapeutic targets after nerve injury.

**Study Design:** Laboratory experiment.

**Methods:** Rat brainstems at E14, E16, E18, E20, adult (4 animals/timepoint) were sectioned and stained for GDNF receptors: GFRα-1, GFRα-2, GFRα-3, and Ret. Islet1 and ChAT were used as markers for motoneuron cell bodies. Sections were observed using Zeiss Axio Imager M2 Microscope and quantified using Image J.

**Results:** Expression of all four GDNF receptors was identified within the nucleus ambiguus, as well as hypoglossal and facial nuclei of the adult rat brainstem. During rat development, GFRα-3 and Ret exhibited upregulation within the nucleus ambiguus at E14 whereas GFRα-1 began showing upregulation at E20. GFRα-2 exhibited no upregulation at embryonic timepoints. Conclusion: Upregulation of the GDNF receptors within the nucleus ambiguus occur after laryngeal muscles innervation during development and may associated with maturation and maintenance of the neuromuscular synapses of the larynx before and after birth. No differences among ambiguus, hypoglossal, and facial nuclei was observed.

## INTRODUCTION

Transection of the recurrent laryngeal nerve (RLN) is a risk of surgeries involving neck or upper thorax pathology, such as thyroidectomies or cervical spine surgeries.^1–3^ RLN injury causes ipsilateral vocal fold paralysis resulting in dysphagia, respiratory distress and dysphonia.^1,4^ Nerve injury leads to Wallerian degeneration of the distal segment of the RLN and regenerated axonal outgrowth from the proximal end of the RLN towards the intrinsic laryngeal muscles (ILM) of the larynx. However, motor reinnervation of the larynx is non-selective.^5^ As a consequence, synkinetic vocal fold movement of the aberrant reinnervation of the larynx persist and the normal motion is never restored.^4^

Synkinetic reinnervation of the larynx is guided by upregulation of different cues triggered by the denervation of the ILM.^6–9^ Neurotrophic factors are proteins whose expression promotes the differentiation, maturation, survival, and maintenance of motoneurons during development and postnatal life.^6–9^ They may also be expressed in denervated muscles, serving as long distance signaling cues for axon regeneration, as has been observed in RLN injury and ILM reinnervation.^10^

Glial Cell-Derived Neurotrophic Factor, GDNF, is a neurotrophic factor originally isolated from rat glial cell lines.^5,11–13^ It is a member of the transforming growth factor-β super family, which activates intracellular signaling for axonal outgrowth and neuronal survival via the receptor tyrosine kinase, Ret. Ret is activated via ligand binding to GPI-linked co-receptors of the GDNF Family Receptor Alpha family.^5,6,8,9,11,12,14^ GDNF and its receptors, Ret and GFRalpha-1/2/3 are expressed within brainstem neurons. Changes in expression of Ret and GFRalpha-1/2/3 were observed during development, in adulthood and post RLN injury in the rat and mouse brainstem.^9^ In the periphery, GDNF and its receptors play a role in neuromuscular junction formation as differential expression of these receptors activates intracellular signaling of pathways responsible for successful axonal guidance and complete innervation of muscle targets.^11,12,14,15^ The motoneurons of the nucleus ambiguus (NAmb) is one such location where GDNF may play roles in axonal guidance and survival properties.^9,16^

NAmb is a long rostrocaudally oriented column of motoneurons located in the ventral aspect of the lower medulla oblongata that innervate structures within the head, neck, thoracic and abdominal cavity.^6,9,11,12,17,18^ The column of NAmb is constituted in three parts, being the ILM motoneurons located in the caudal third of the column supplying the larynx by the superior laryngeal nerve and the RLN.^18–22^ The origin of NAmb begins during medulla oblongata formation at day 12 of rat development.^23^ GFRalpha-1/2/3 and Ret have shown immunoreactivity at cranial nuclei in the medulla in formation, as well as within neuronal cell populations of the NAmb, suggesting that the GDNF receptors play an appreciable role during and after development.^9^

While expression of GDNF and its receptors have been studied in murine and human brainstem development, differential expression of GDNF receptors at chronological developmental stages, in relation to ILM innervation, is a novel area of study as it pertains to recurrent laryngeal nerve injury.^9^ Thus, our study aims to elucidate individual roles of GDNF receptors, GFRalpha1/2/3 and Ret within NAmb to determine their role in the innervation of the ILM during laryngeal embryogenesis. Proper identification of GDNF receptor expression in the adult and embryonic brainstem will aid in subsequent development of a somatotrophic map along NAmb.

## MATERIAL AND METHODS

### Animals

During this study, regulations and laws regarding animal handling and care were upheld in accordance with the protocols of the Institutional Animal Care and Use Committee with training conducted at the Institute of Comparative Medicine and lab supervision under Columbia University Environmental Health & Safety. Four adults and embryonic 32 rats of different development stages were analyzed in the present work: E14, E16, E18 and E20 days (4 animals per group). All adult rats and litter mothers were sacrificed via intraperitoneal injection of xylazine (20 mg/ml) and ketamine hydrochloride (100 mg/mL). Successful euthanasia was confirmed by assessment of respiratory effort and hindlimb toe pinch.

### Tissue Extraction

Following euthanasia, adult animals underwent trans-cardiac perfusion with 0.1 M phosphate buffer saline (PBS) (200 ml) followed by 4% paraformaldehyde in PBS (250 ml). To access the brainstem the skull was removed, and the rat brainstem was dissected caudally at the spinomedullary junction and rostrally at the junction of the midbrain and diencephalon. Brainstems were post-fixed in vials of 4% paraformaldehyde in PBS at 4 degrees overnight. The brainstems were immersed in 15% sucrose in PBS until the tissue sunk. The pieces were transferred to 30% sucrose in PBS until sunk prior to be being submerged in cryostat embedding medium in a mold and frozen at -80C.

In gestational stages, embryos were extracted from euthanized pregnant dams at different developmental periods as mentioned above. After confirmation of euthanasia, a vertical midline incision through the abdomen was performed using a scalpel. Flaps were reflected and pinned bilaterally as to expose the gravid uterus. Embryonic yolk sacs were carefully punctured and peeled until embryos could be lightly squeezed out of their sac’s. Embryos were placed in vials of 4% paraformaldehyde in PBS at 4° C overnight. Embryos were then submerged in sucrose 15% and 30% prior to processing. Embryos was dissected at the base of the neck. The piece was then positioned at a 50°-60° angle in relation to the horizontal axis of the mold, covered with cryostat embedding medium, and frozen at -80C.

### Sectioning

Molds were placed in a -20° C cryostat for 30 minutes. Adult brainstems were sectioned in coronal sections at 40 µm caudal to rostral and placed serially in twenty-four multiwell plates containing PBS. Embryos were sectioned in coronal sections from caudal level at the area of the larynx and cervical spinal cord to cranial level at the cerebellar pontine angle in the rostral hindbrain. All embryos were sectioned at 14 µm. Coronal sections of the embryos were placed onto gelatin slides and left to dry prior to immunostaining.

### Immunohistochemistry

Two separate IHC protocols were utilized for this study. The IHC protocol was carried out in floating sections for adult brainstems and sections on gelatin slides for the embryos. Brainstems were stained for motoneuron marker Islet-1 (39.4DS, Developmental Systems Hybridoma Bank, Iowa City, Iowa) and for GDNF Receptors Ret, GFRα-1, GFRα-2, and GFRα-3.

Adult brainstem sections were kept in twenty-four multiwell plates. They were divided into six columns. Each column was blocked for 30 minutes with 1% donkey serum in 0.3% triton in PBS (PBST) and then incubated in 1% donkey serum Goat in PBST with anti-ChAT antibody plus another primary antibody for GDNF receptors as summarized in Table 1. The sections were incubated, on a BioRocker 2D shaker, in primary antibody solutions for 72 hours at 4° C. Sections were washed twice with PBS for 5 minutes prior to incubation in the secondary antibody solutions for two hours as stated in Table 1. Floating sections were captured onto gelatin slides in serial order from caudal to rostral brainstem and mounted with 4’6’-diamidino-2-phenylindole dihydrochloride (DAPI) (Abcam, Cambridge, MA).

**Table 1.** Number of animals used in this study

| Experiments | Groups |  |  |  |  | Post-fixation | Section thickness | Immunohistochemistry incubation |  |
| --- | --- | --- | --- | --- | --- | --- | --- | --- | --- |
|  | E14 | E16 | E18 | E20 | Adult |  |  | Primary antibody | Secondary antibody |
| Islet1/GFRα1/DAPI | 4 | 2 | 4 | 2 | | 48 hours | 14 μm | Mouse anti-Islet 1 [39.4D5; DSHB: Iowa City, Iowa] (1:50) & Goat anti-GFRα-1 R&D Systems; Minneapolis, MN (1:100) | Ab2: DaM-AF488 [Invitrogen: Waltham, MA] (1:200) + DaG-AF594 [Invitrogen: Waltham, MA] (1:200) |
| Islet1/GFRα2/DAPI | 4 | 3 | 4 | 1 | | 48 hours | 14 μm | Mouse anti-Islet 1 [39.4D5; DSHB: Iowa City, Iowa] (1:50) & Goat anti-GFRα-2[R&D Systems; Minneapolis, MN] (1:100) | Ab2: DaM-AF488 [Invitrogen: Waltham, MA] (1:200) + DaG-AF594 [Invitrogen: Waltham, MA] (1:200) |
| Islet1/GFRα3/DAPI | 4 | 2 | 4 | 2 | | 48 hours | 14 μm | Mouse anti-Islet 1 [39.4D5; DSHB: Iowa City, Iowa] (1:50) & Goat anti-GFRα-3[R&D Systems; Minneapolis, MN] (1:100) | Ab2: DaM-AF488 [Invitrogen: Waltham, MA] (1:200) + DaG-AF594 [Invitrogen: Waltham, MA] (1:200) |
| Islet1/Ret/DAPI | | | | | 4 | 48 hours | 14 μm | Mouse anti-Islet 1 [39.4D5; DSHB: Iowa City, Iowa] (1:50) & Goat anti-Ret [R&D Systems; Minneapolis, MN] (1:100) | Ab2: DaM-AF488 [Invitrogen: Waltham, MA] (1:200) + DaG-AF594 [Invitrogen: Waltham, MA] (1:200) |
| ChAT/DAPI | | | | | 3 | 72 hours | 40 μm | Ab1: Goat anti-ChAT (1:100) | DaG-AF594 (1:200) |

GFRα : GDNF Family Receptor Alpha; Ret: Rearranged during Transfection gene receptor; DAPI: 4',6-diamidino-2-phenylindole; ChAT: Anti-Acetylcholine Transferase; DaM-AF488: Donkey anti-Mouse conjugated to Alexa Fluor 488; DaG-AF594: Donkey anti-Goat conjugated to Alexa Fluor 594; DaG-AF594: Donkey anti-Goat conjugated to Alexa Fluor 594

For the embryological groups, sections were placed on gelatin slides. ImmEdge™ Pen (Burlingame, CA) was used for blocking the staining area and slides were placed in a slide box to dry. Sections were postfixed in 4% paraformaldehyde in PBS for 10 minutes at room temperature followed by a wash with PBS for 5 minutes. Antigen retrieval was performed via incubation in citrate buffer solution 1% 1mM (pH=6) for 45 minutes at 60° C. After a wash, sections were blocked as stated above before incubation with the primary antibody solution consisting of Islet 1 plus another primary marker as stated in Table 1. Slides were incubated in the 4° C refrigerator for 48-72 hours. After two washes for 5 minutes in PBS sections were incubated in secondary antibody solution for 2 hours at room temperature. Slides were mounted in Flouroshield mounting Medium with DAPI.

### Analysis

Slides were observed at 20X and 40X, and images taken using Zeiss LSM Confocal and Zeiss Axio Imager M2 epi-fluorescences microscopes (Zeiss, Oberkochen, Germany). Images taken were then analyzed using Fiji software to evaluate the timing and level of upregulation of receptors in the motoneurons of the nucleus ambiguus and other nuclei of the brainstem. Positive cell bodies for positive for Islet 1 and receptors were quantified through manual counting under the microscope. Descriptive statistics such as mean, median, and interquartile range from cell counting were processed using R studio application. ANOVA and t test statistical analysis was used to compare variance among the timepoints and receptor populations.

## RESULTS

In this study immunoreactivity for GDNF receptors, GFRalpha-1/2/3 and Ret was observed within the NAmb, hypoglossal, and facial nuclei of the adult brainstem. We also observed GFRalpha-3 and Ret immunoreactivity within the NAmb as early as day 14 of development, whereas GFRalpha-1 was undetectable in the NAmb until 16 days. GFRalpha-2 showed no upregulation in NAmb from 14 to 20 developmental days (Figure 2 and Table 2).

**Table 2.**
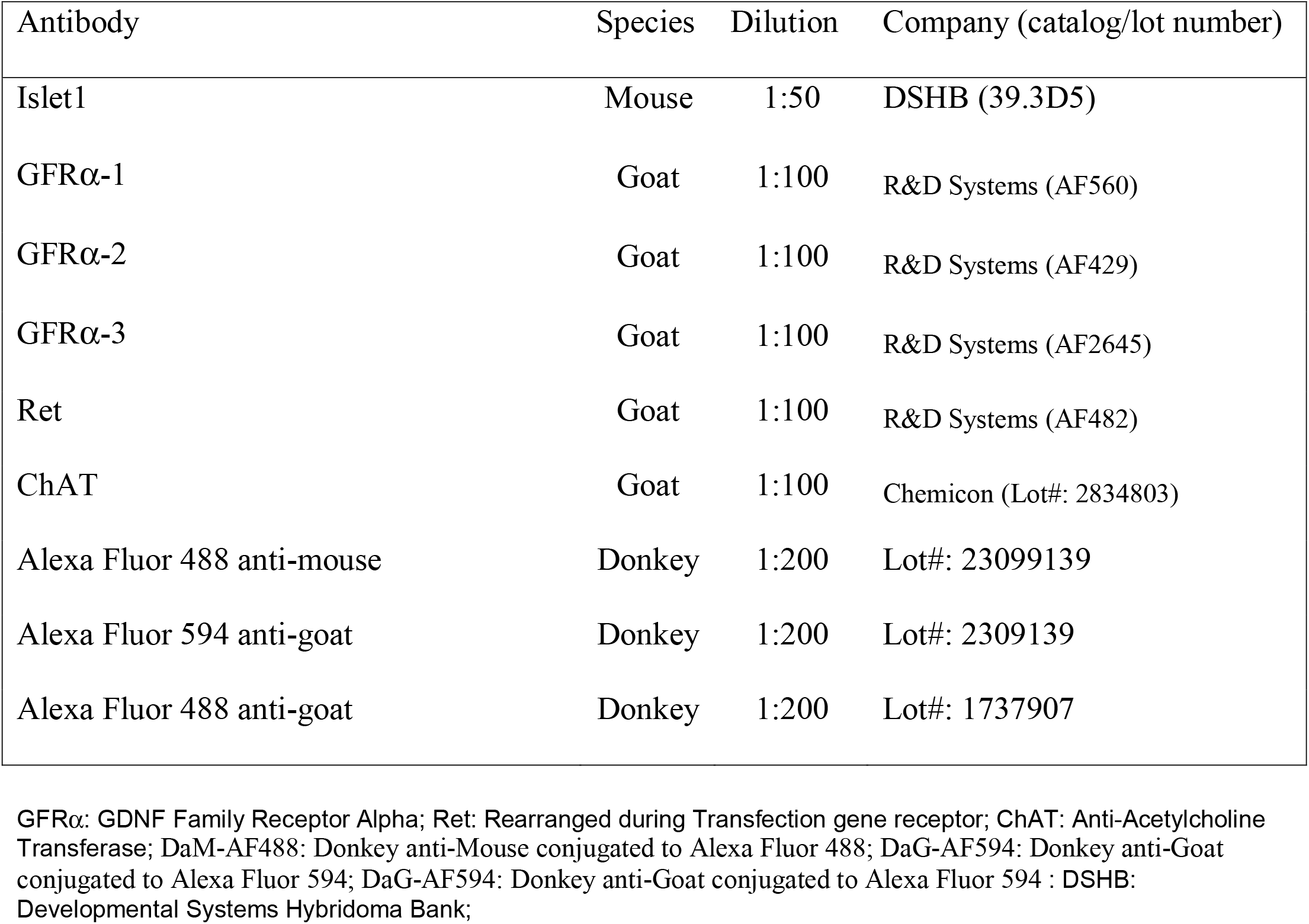
Summary of GDNF Receptor expression at within the motoneurons and nerve fibers of the Nucleus Ambiguus in adult brainstem as well as E14, E16, E18, and E20 of development. (-) negative signal; (-/+) variable negative/weak positive signal; (+) positive signal.

| Antibody | Species | Dilution | Company (catalog/lot number) |
| --- | --- | --- | --- |
| Islet1 | Mouse | 1:50 | DSHB (39.3D5) |
| GFRα-1 | Goat | 1:100 | R&D Systems (AF560) |
| GFRα-2 | Goat | 1:100 | R&D Systems (AF429) |
| GFRα-3 | Goat | 1:100 | R&D Systems (AF2645) |
| Ret | Goat | 1:100 | R&D Systems (AF482) |
| ChAT | Goat | 1:100 | Chemicon (Lot#: 2834803) |
| Alexa Fluor 488 anti-mouse | Donkey | 1:200 | Lot#: 23099139 |
| Alexa Fluor 594 anti-goat | Donkey | 1:200 | Lot#: 2309139 |
| Alexa Fluor 488 anti-goat | Donkey | 1:200 | Lot#: 1737907 |

GFRα : GDNF Family Receptor Alpha; Ret: Rearranged during Transfection gene receptor; ChAT: Anti-Acetylcholine Transferase; DaM-AF488: Donkey anti-Mouse conjugated to Alexa Fluor 488; DaG-AF594: Donkey anti-Goat conjugated to Alexa Fluor 594; DaG-AF594: Donkey anti-Goat conjugated to Alexa Fluor 594 : DSHB: Developmental Systems Hybridoma Bank;

To identify the different motoneuron pools of the brainstem we used Islet1 and Choline acetyltransferase (ChAT) in adult animals. ChAT was clearly identified in the cytoplasm of the motoneurons that constitutes the adult NAmb, hypoglossal, and facial nuclei of the medulla (Figure 1). Islet1 and ChAT immunoreactive neurons were also observed in those that constitute the oculomotor nuclei within caudal levels of the Pons. While there was consistent intensity of Islet1 and ChAT within the hypoglossal, facial and oculomotor nuclei, the NAmb displayed varying levels of upregulation among the cell bodies of the adult NAmb. Quantification of cells within the NAmb yielded a mean of 645 ± 141 (median = 606) immunoreactive motoneurons.

**Fig. 1.**
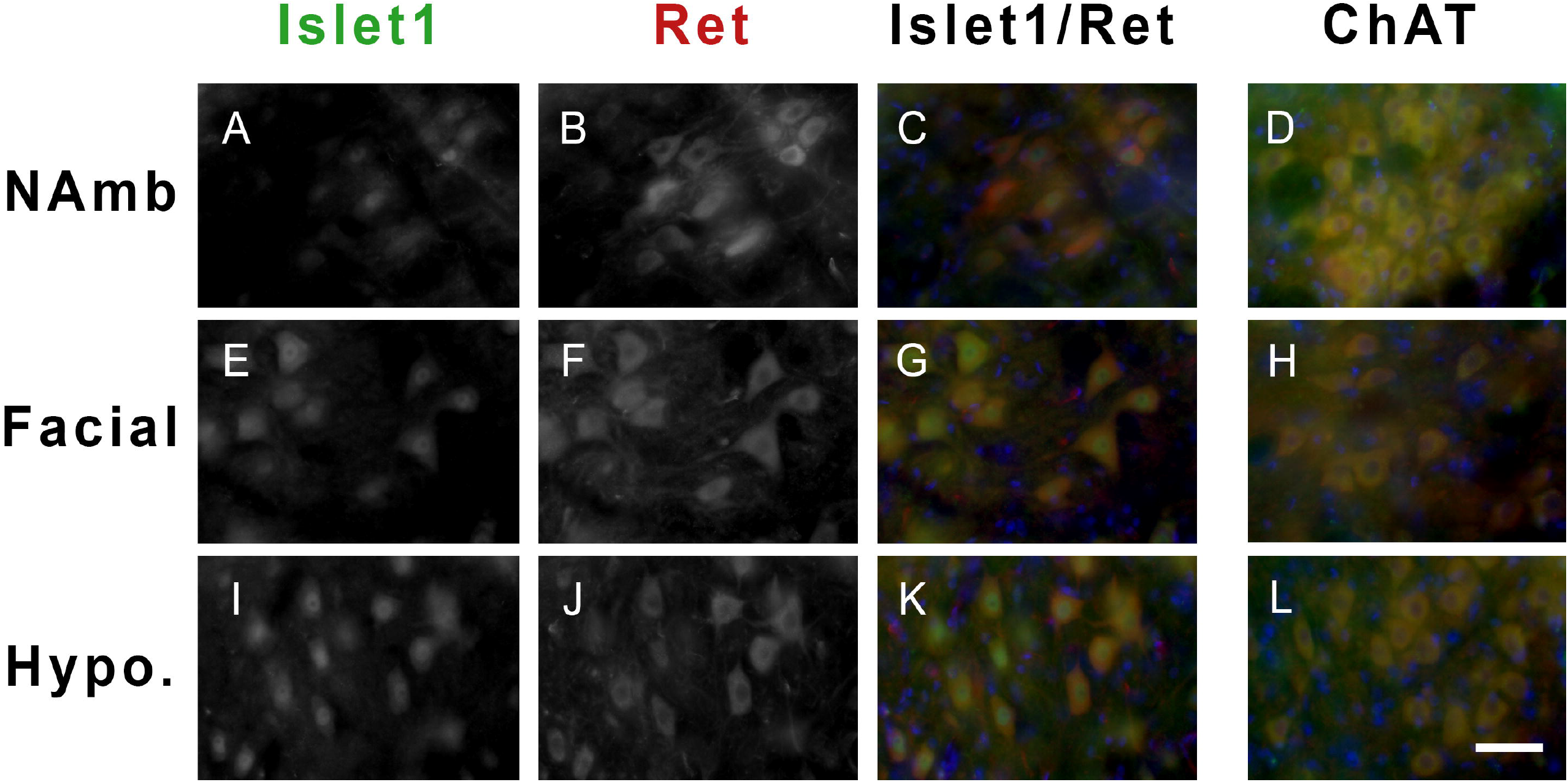
Staining of Adult Nucleus Ambiguus (A-D), Facial nucleus (E-H), and Hypoglossal nucleus (I-L). Islet1 labeling of the adult NAmb (A), Facial nucleus (E), and Hypoglossal nucleus (I). Labeling with Ret within the adult NAmb (B), Facial nucleus (F), and Hypoglossal nucleus (J). Merged image of Islet1 (green) and Ret (red) in the adult NAmb (C), Facial nucleus (G), and Hypoglossal nucleus (K). Merged imaged of ChAT (Acetylcholine Transferase) in red and Oct6 (green) within the adult NAmb (D), Facial nucleus (H), and Hypoglossal nucleus (L). Scale bar = 50 µm

**Fig. 2.**
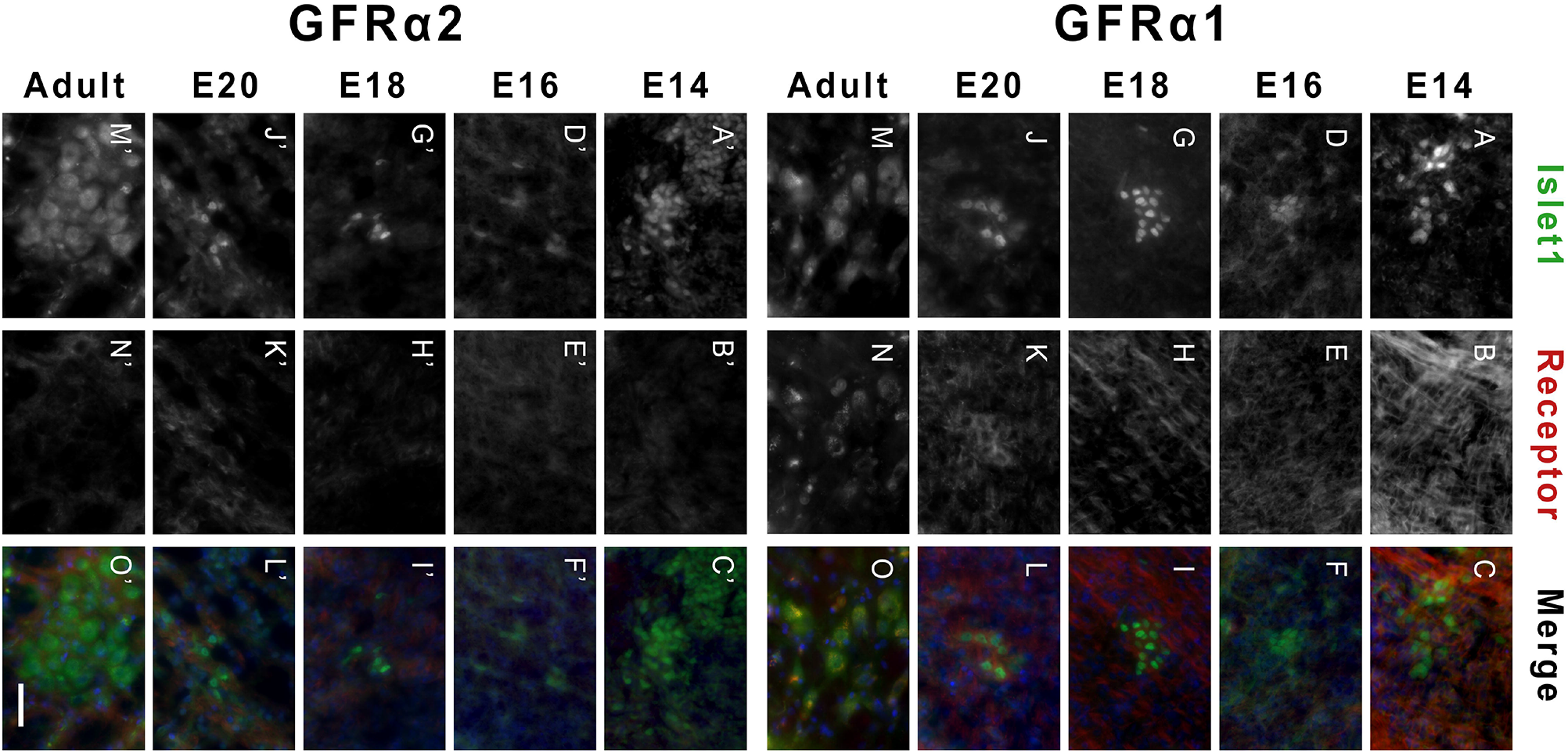
Labeling of nucleus ambiguus motoneurons in adult and E14-E20 embryos with Islet1 (A/A’, D/D’, G/G’, J/J’, M/M’). Labeling of nucleus ambiguus motoneurons and/or nerve fibers with GFR⍰1 (B, E, H, K, N). Labeling of nucleus ambiguus motoneurons and/or nerve fibers with GFR⍰ 2 (B’, E’, H’, K’, N’). Merged image of Islet1 and GFR⍰ 1 (C, F, I, L, O). Merged image of Islet1 and GFR⍰ 2 (C’, F’, I’, L’, O’). Scale bar = 50 µm

In developmental stages, differential Islet1 labelling was identified along studied timepoints. Namely, Islet-1 expression was observed within the nucleus ambiguus, hypoglossal nuclei, facial nerve, and nodose ganglia as well as developing ILM (Figure 2 and 3). Despite clear labelling of Islet1 allowing for unambiguous identification of the NAmb, a different intensity was observed when compared to the other motor nuclei of the brainstem such as facial or hyoglossus nuclei. Those nuclei and nodose appear brighter than motoneurons of the NAmb at all embryonic timepoints (Figure 2 and 3). The adult brainstem displayed a moderate signal in the cell bodies of the NAmb and strong signal in the hypoglossal and facial nuclei (Figure 1).

**Fig. 3.**
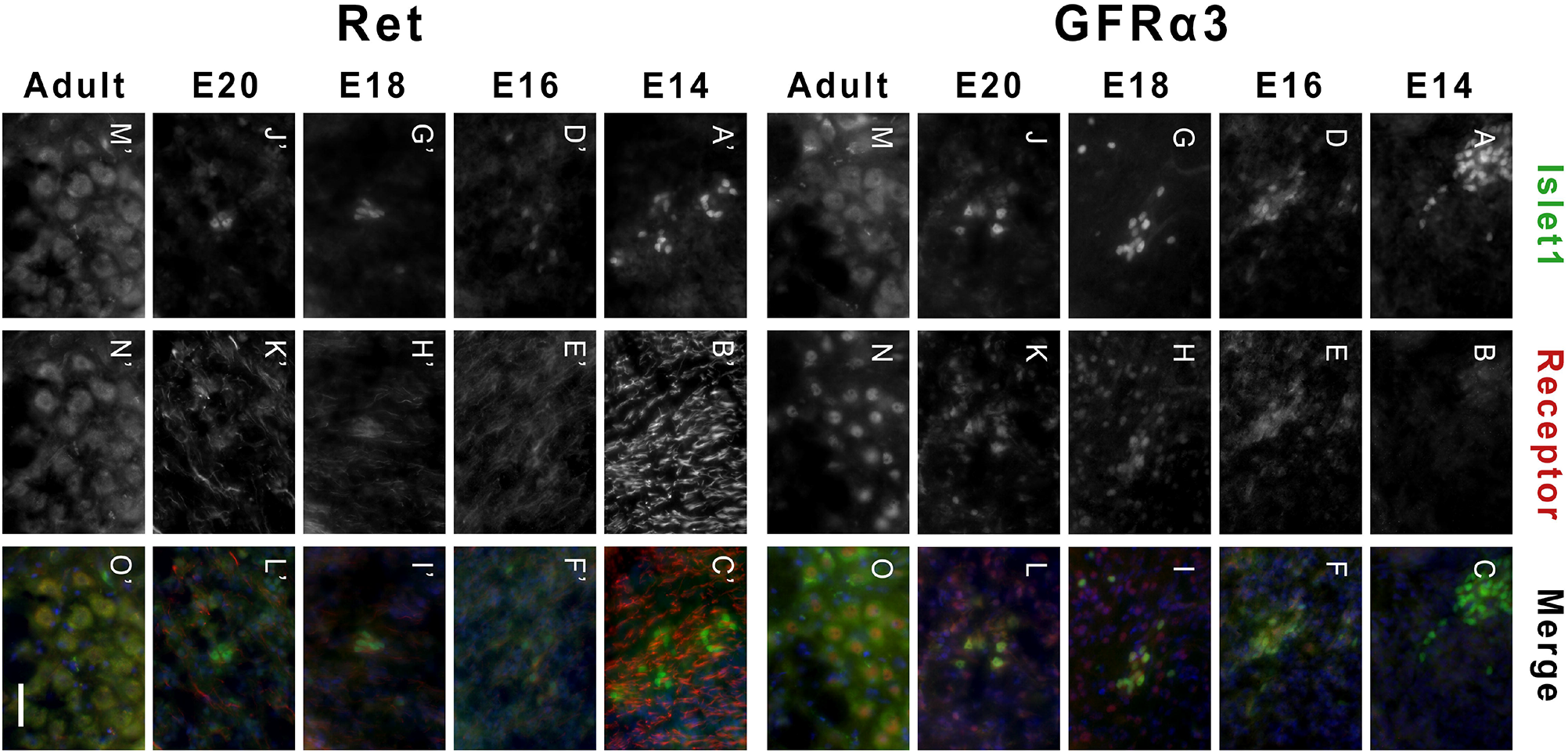
Labeling of nucleus ambiguus motoneurons in adult and E14-E20 embryos with Islet1 (A/A’, D/D’, G/G’, J/J’, M/M’). Labeling of nucleus ambiguus motoneurons and/or nerve fibers with GFR⍰ 3 (B, E, H, K, N). Labeling of nucleus ambiguus motoneurons and/or nerve fibers with Ret (B’, E’, H’, K’, N’). Merged image of Islet1 and GFR⍰ 3 (C, F, I, L, O). Merged image of Islet1 and Ret (C’, F’, I’, L’, O’). Scale bar = 50 µm

GFRalpha-1 positive staining was observed in NAmb neurons in the adult brainstem (94 ± 74 (median = 109) neurons positive for GFRalpha-1 at adult Namb, Table 3). At E14, GFRalpha-1 showed upregulation in the nerve fibers at ventromedial aspect of the brainstem yet the dorsolateral aspect and individual neurons remained negative. At E16, a positive staining for GFRalpha-1 staining was showed in the nerve fibers run along a ventromedial course that is highly specific in pattern and direction (Figure 4). A positive staining was also observed in the intrinsic muscles of the larynx at this timepoint. As the rat brainstem progresses to E18, an increase in GFRalpha-1 signal was specifically observed at the spinal trigeminal tract and the nerve fibers forming cuneatus and fragilis tracts at the hindbrain, while the Namb, hypoglossus and facial nuclei remain negative (Figure 2). Of note, while the hypoglossal appears negative internally, there is distinct uptake observed around the external lining of the hypoglossal cells in what appears to be a honeycomb presentation. No changes were appreciated between E18 and E20 brainstems.

**Table 3.** Data Summary – Qualitative

|  | **GFRα1**<br>**Neurons** | **GFRα1**<br>**Nerve**<br>**Fibers** | **GFRα2**<br>**Neurons** | **GFRα2**<br>**Nerve**<br>**Fibers** | **GFRα3**<br>**Neurons** | **GFRα3**<br>**Nerve**<br>**Fibers** | **Ret**<br>**Neurons** | **Ret**<br>**Nerve**<br>**Fibers** |
| --- | --- | --- | --- | --- | --- | --- | --- | --- |
| **E14.5** | - | + | - | - | - | - | - | + |
| **E16.5** | - | + | - | - | + | - | - | + |
| **E18.5** | -/+ | + | - | + | + | - | - | + |
| **E20.5** | + | + | - | + | + | - | - | + |
| **Adult** | + | + | - | + | + | - | + | + |

**Table 4.**
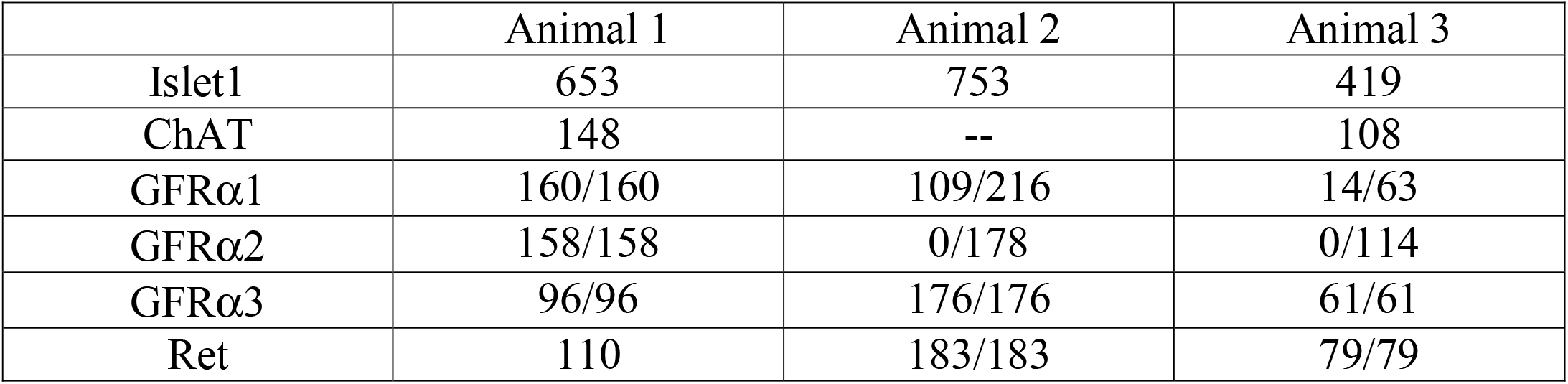
Quantification of Nucleus Ambiguus Motoneurons within the Adult Brainstem

**Fig. 4.**
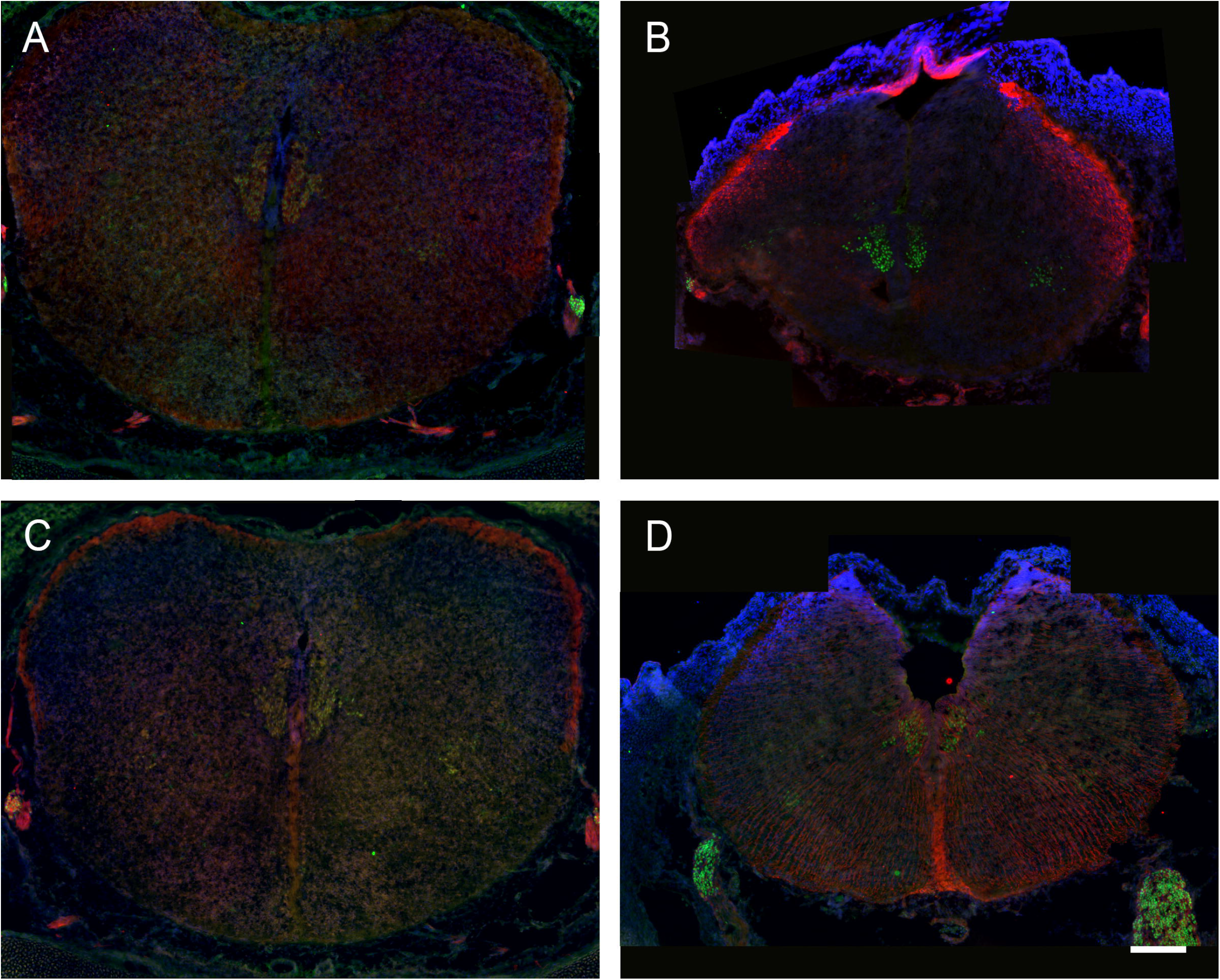
Rat medulla oblongata at E16. Labeling of motoneurons of the nucleus ambiguus and hypoglossal nucleus in green and nerve fibers in red (A-D). Labeling of nuclei of the nucleus ambiguus, hypoglossal, and interneurons with GFRalpha-3 in red (C). GFRalpha-1 in red (A); GFRalpha-2 in red (B); GFRalpha-3 in red (C); Ret in red (D) Scale bar = 200 µm

Adult brainstem showed variability in which three of four specimens were negative for GFRalpha-2 in all anatomical regions, whereas one specimen showed immunoreactivity for the NAmb, facial, oculomotor, and hypoglossal nuclei (Figure 5). At E14, GFRalpha-2 staining was negative in the hindbrain except for a weak signal at the spinal trigeminal tract in the dorsolateral aspect of the brainstem in formation. As the hindbrain develops, at E16, a stronger intensity was noted at the spinal trigeminal tracts nerve fibers. In contrast, a faint immunoreactivity was observed in the rest of the hindbrain areas and the larynx (Figure 2 and 4). AT E18, an appreciable increase of intensity of GFRalpha-2 staining was identified in the ventromedial aspect of the hindbrain, while immunoreactivity of the motor nuclei of the medulla was negligible. No changes were identified in the E20 brainstem (Figure 2).

**Fig. 5.**
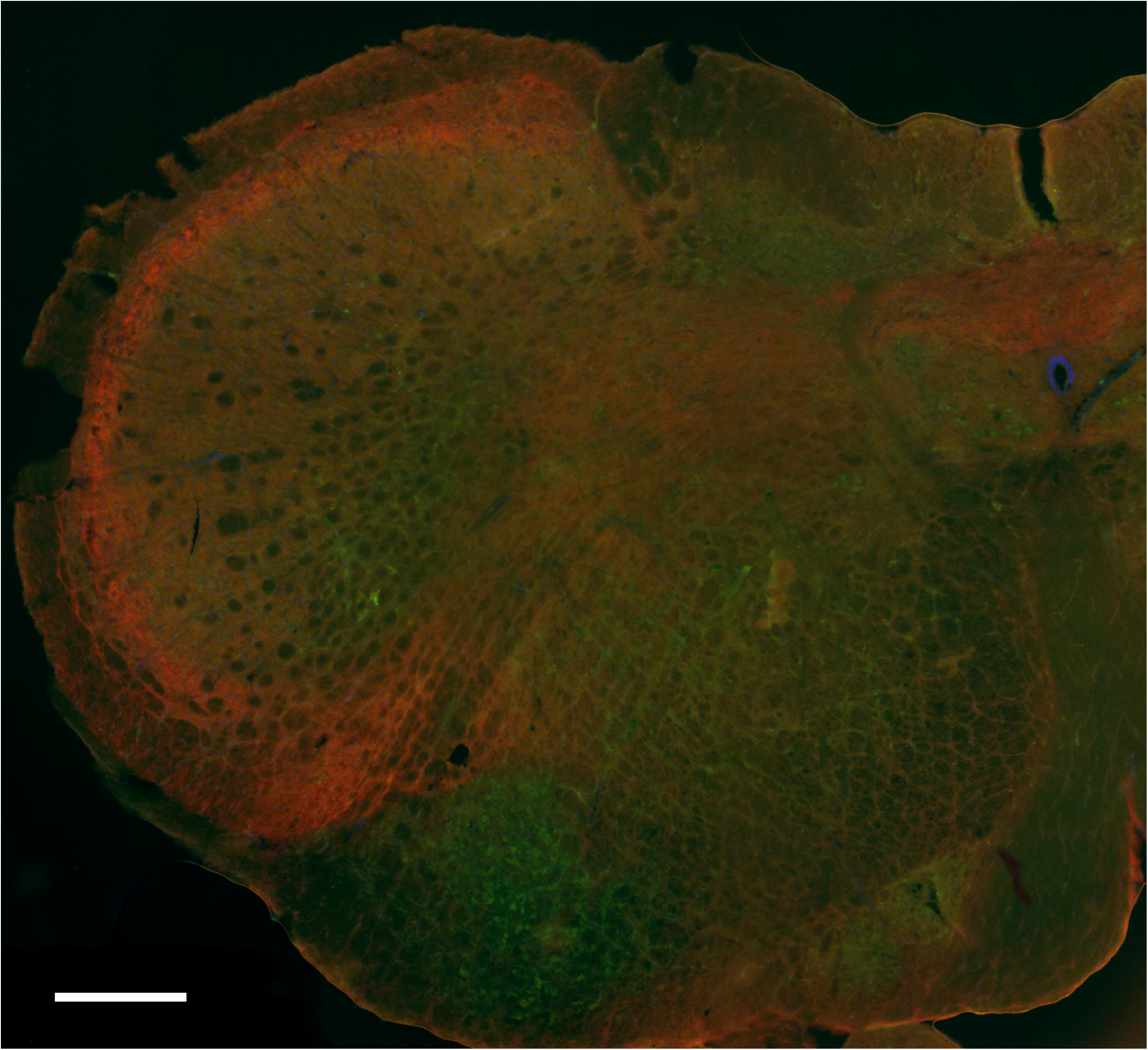
Hemisection of adult medulla oblongata with motoneuron labeling with Islet1 in green and labeling of nerve fibers with GFRalpha-2 in red. Scale bar = 200 µm

In adult brainstems no differences were identified regarding GFRalpha-3 immunoreactivity among any region of the medulla oblongata. On average, there were 111 ± 59 (median = 96) GFRalpha-3 positive motoneurons within the NAmb (Table 3). In E14 rat hindbrains, GFRalpha-3 was undetectable in all studied embryos (Figure 3). In contrast, the nodose ganglia displayed strong immunoreactivity within its nerve fibers. As the hindbrain progressed to E16, an increase of the GFRalpha-3 intensity was identified in the spinal trigeminal tract yet negative staining observed in motor nuclei of the medulla in formation. outside of the nervous system, weak staining was observed and the mucosa of the pharyx and the larynx. The most dramatic change in GFRalpha-3 labelling happened at E18 in the hindbrain, where global nuclear immunoreactivty throughout the entirety of the rat medulla was observed.

Positive immunoreactivity for Ret was observed initially in the adult brainstem in all animals (Figure 3). Ret was positive within cell bodies of the NAmb, facial, hypoglossal, and oculomotor nerve. Moreover, nerve fiber also exhibited immunoreactivity for GFRalpha-3 albeit weaker in adult medulla compared to the embryos. Of the three nuclei at the medulla, Ret immunoreactivity was weakest in NAmb while the other facial and hypoglossus nuclei displayed stronger labelling. On average, there were 124 ± 53 (median = 110) Ret positive motoneurons within the adult NAmb. At E14, Ret was strongly immunopositive signal within nerve fibers of the hindbrain running in a distinct ventromedial direction. Ret immunoreactivity was also observed at the nodose ganglia. This pattern of immunoreactivity remained through development of the E16 hindbrain but a weaker labelling for Ret was observed within the nodose ganglia. No differences were observed at E18 or E20 group except for an increased intensity within all muscles of the head, including those of the ILM.

## DISCUSSION

Results of this study showed GDNF receptors production during development within hindbrain and suggest they play a role in development and maintenance of the rat NAmb.

Islet1 labelling within the Namb was variable among embryonic timepoints. Namely there was an apparent decrease of Islet1 signal from E14 to E16 followed by stronger immunoreactivity from E16 to E18. Adult brainstems also showed inconsistency regarding Islet1 at NAmb and in comparation to different nuclei of the medulla oblongata. On the other hand, the facial and hypoglossal nuclei expressed Islet1 at an apparent higher intensity than the observed in NAmb. A possible explanation may lie due to the orientation of the NAmb and the remaining nuclei of the medulla. NAmb constitutes a rostrocaudal column of motoneurons, in contrast to the mediolateral axis of the facial nucleus and the dorsoventral axis of the hypoglossal nucleus. Another explanation for this inconsistency could be inherent differences regarding Islet1 regulation in the different motoneuron pools within brainstem. Further experiments exploring colocalization of ChAT with Islet1 in embryonic brainstems may be useful in understanding Islet1 role in NAmb during development.

GDNF receptors were all positive within the Namb, hypoglossal, and facial nucleus except for GFRalpha-2 (Figure 2 and 3). Labelling was consistent with a prior research which found positive upregulations of all GDNF receptors within the ILM following RLN injury, with the exception of GFRalpha-2.^11,16,9^ Moreover, the quantification of NAmb motoneurons was consistent with studies found in the literature regarding location, extension and number of motoneurons identified. ^18^

The neurons forming NAmb were identified at E14 and beyond. In contrast, a differential expression of GDNF receptors at all time points and adulthood was observed. Immunoreactivity for GFRalpha-1, GFRalpha-3, and Ret was observed during development at different developmental times and continued into adulthood. In contrast, GFRalpha-2 immunoreactivity was undetectable at all embryonic timepoints yet its labelling within nerve fibers of the medulla oblongata suggest an undetermined role in axonal development.

## CONCLUSION

Receptor expression during development is chronological though all receptors exhibit a different pattern of localization. Colocalization of Islet1 with GDNF receptors showed differences in NAmb compared to other nuclei in the medulla oblongata. Further research on this subject may be useful in providing insight to therapeutic advancements to vocal cord pathology following RLN injury.

## Notes

**SOURCES OF FUDING:** This work was supported by the National Institutes of Health: 1R01DC018060

### Competing Interest Statement

The authors have declared no competing interest.

